# Patterns of divergence across the geographic and genomic landscape of a butterfly hybrid zone associated with a climatic gradient

**DOI:** 10.1101/149179

**Authors:** Sean F. Ryan, Michael C. Fontaine, J. Mark Scriber, Michael E. Pfrender, Shawn T. O’Neil, Jessica J. Hellmann

## Abstract

Hybrid zones are a valuable tool for studying the process of speciation and for identifying the genomic regions undergoing divergence and the ecological (extrinsic) and non-ecological (intrinsic) factors involved. Here, we explored the genomic and geographic landscape of divergence in a hybrid zone between *Papilio glaucus* and *Papilio canadensis*. Using a genome scan of 28,417 ddRAD SNPs, we identified genomic regions under possible selection and examined their distribution in the context of previously identified candidate genes for ecological adaptations. We showed that differentiation was genome-wide, including multiple candidate genes for ecological adaptations, particularly those involved in seasonal adaptation and host plant detoxification. The Z-chromosome and four autosomes showed a disproportionate amount of differentiation, suggesting genes on these chromosomes play a potential role in reproductive isolation. Cline analyses of significantly differentiated genomic SNPs, and of species diagnostic genetic markers, showed a high degree of geographic coincidence (81%) and concordance (80%) and were associated with the geographic distribution of a climate-mediated developmental threshold (length of the growing season). A relatively large proportion (1.3%) of the outliers for divergent selection were not associated with candidate genes for ecological adaptations and may reflect the presence of previously unrecognized intrinsic barriers between these species. These results suggest that exogenous (climate-mediated) and endogenous (unknown) clines may have become coupled and act together to reinforce reproductive isolation. This approach of assessing divergence across both the genomic and geographic landscape can provide insight about the interplay between the genetic architecture of reproductive isolation and endogenous and exogenous selection.

## Introduction

Hybrid zones are ideal systems for exploring the interplay between exogenous (ecological factors, such as climate) and endogenous (genetic incompatibilities) selective pressures that drive genomic divergence. Although climate has been implicated in the maintenance of a number of hybrid zones across a broad range of taxa (Taylor *et al*. 2015), our understanding of how climate drives patterns of divergence and gene flow between the genomes of hybridizing species remains limited to a handful of systems (Taylor *et al*. 2014; de Villemereuil *et al*. 2014; Hamilton *et al*. 2014).

It has long been recognized that an advantage to studying hybrid zones is their utility in understanding the genetic basis of species divergence, i.e., identifying those regions of the genome under selection and potentially involved in reproductive isolation (Barton & Hewitt 1985). A classical and still widely used approach is to look at how genetic differences between species vary across geography and species boundaries, through a geographic cline analysis. Complementary to cline analysis, there are a number of population genomic approaches to identify genomic regions under selection (i.e., via “genome scans”) that do not require spatial sampling along a cline. Genome scans can be used to identify outlier loci from a null expectation based on genome-wide ancestry that are potential indicators of divergent selection (Foll & Gaggiotti 2008). Given that geographic cline analyses and genome scans estimate selection dynamics operating on different spatiotemporal scales, from recent to more long-term time scale respectively, combing these approaches increases the power to detect regions of the genome involved in local adaptation and reproductive isolation (de Lafontaine *et al*. 2015).

Placing patterns of genetic differentiation within the genomic landscape can also provide insight as to how genetic architecture itself influences introgression across the genome—the “genomic architecture of divergence” (Teeter *et al*. 2010; Gompert *et al*. 2012). A number of studies suggest that gene flow may be heterogeneous across the genome of hybridizing species (Mallet 2005; Teeter *et al*. 2008), with large amounts of the genome diffusing between species while a few regions maintain exceptionally high levels of divergence (Fontaine *et al*. 2015). Specifically, regions of low recombination such as inversions, centromeres and sex chromosomes are often found to have elevated levels of divergence and often harbor a disproportionate amount of the factors responsible for reproductive isolation (Coyne & Orr 1989; Presgraves 2008).

Both exogenous and endogenous factors can act to inhibit introgression and maintain species boundaries. Endogenous factors have often been found to play a major role in driving clinal patterns of genetic variation across many hybrid zones (Bierne *et al*. 2011). For example, most tension zones, hybrid zones maintained by a balance between selection and migration, are believed to be the result of intrinsic incompatibles. The geographic selection gradient model (GSGM), a mathematically similar model to the tension zone model, can also produce similar clinal patterns, with the main difference being selection is primarily exogenous (e.g., related to abiotic factors) (Endler 1977). However, endogenous and exogenous barriers are not mutually exclusive (Barton & Hewitt 1985; Hewitt 1988; Bierne *et al*. 2011). In fact, these barriers may become “coupled” and act synergistically to increase reproductive isolation and maintain the location of the hybrid zone, often resulting in tension zones becoming associated (co-occurring) within an ecological gradient (Barton & Hewitt 1985; Bierne *et al*. 2011).

The hybrid zone between the eastern tiger swallowtail (*Papilio glaucus*) and the Canadian tiger swallowtail (*P. canadensis*) butterfly is an excellent system for exploring how climate influences genomic patterns of divergence. The ecology of this system has been studied extensively for over thirty years and climate is implicated as a prominent factor in maintaining reproductive isolation between the two species (Scriber 2011). However, an understanding of how endogenous and exogenous selective forces maintain divergence across the genome remains unresolved. The two species are estimated to have diverged 0.5-0.6 million years ago (Cong *et al*. 2015) and previous genetic and morphological analysis of individuals suggest they form a narrow (< 50 km) east-west hybrid zone in the north eastern United States, extending from Minnesota to New England (Fig 1A) (Scriber 1990). A number of phenotypic traits, including those that are physiological (Scriber *et al*. 1989; Kukal *et al*. 1991; Mercader & Scriber 2008), behavioral (Scriber *et al*. 1991; Deering & Scriber 2002; Mercader *et al*. 2009), and morphological (Luebke *et al*. 1988; Lehnert *et al*. 2012), appear to coincide (exhibit similar clines) with latitudinal and altitudinal variation in climate (Hagen & Scriber 1989; Scriber 2002; Scriber *et al*. 2003). A better understanding of how genes putatively involved in ecological adaptations in this system covary with environmental factors (the geographic landscape) and how they are distributed across the genome (the genomic landscape) will help to elucidate how ecological selection pressures drive reproductive isolation within the genomes of these hybridizing species.

**Figure 1:**
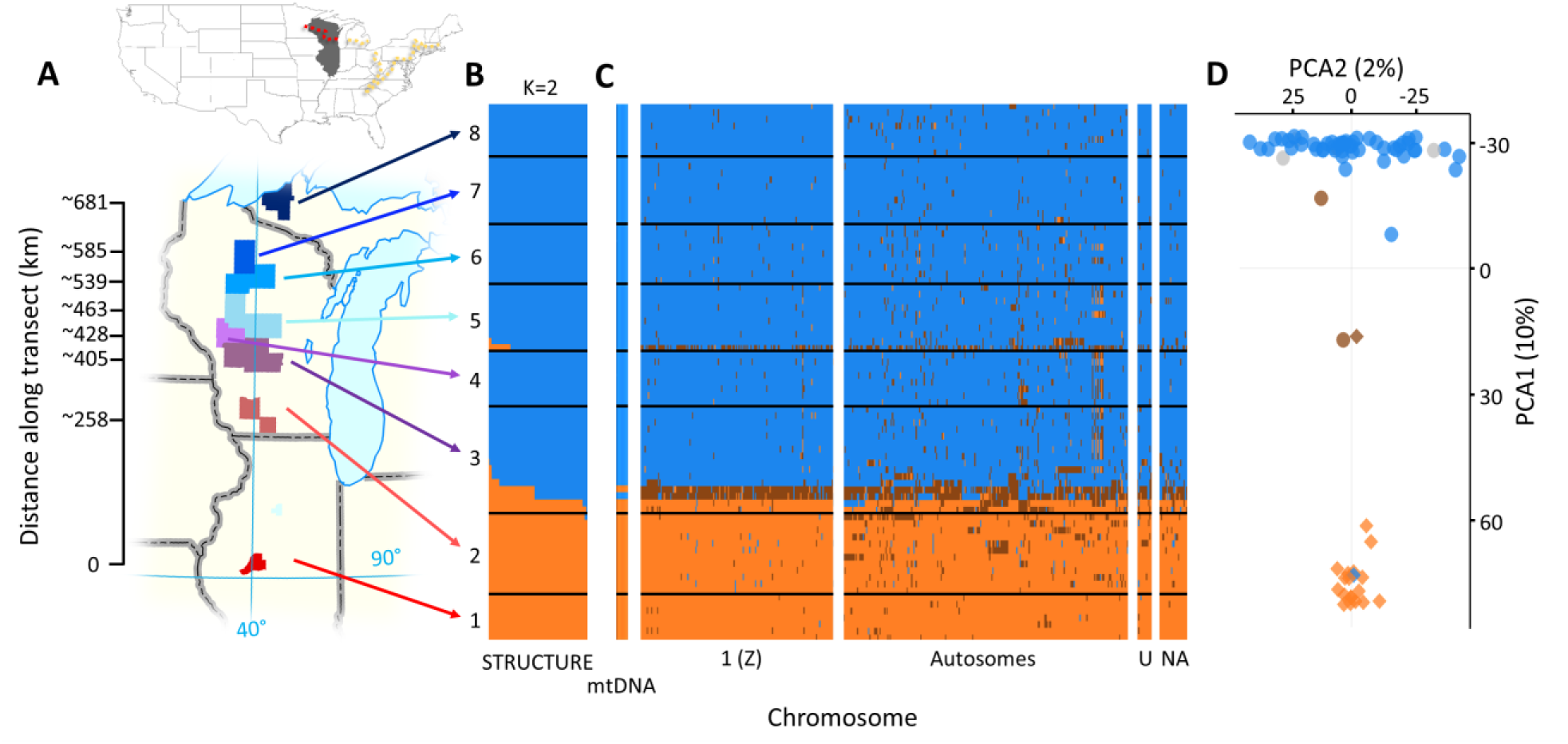
The genomic landscape of divergence: evidence of ecological selection across the genome. (A) Map showing sampling locations for each population; counties with the same color were pooled for analysis. (B) Bayesian clustering of ancestry proportion to K==. (C) Maximum likelihood estimates of hybrid index of one mtDNA marker and 471 SNPs. Each row is an individual and column a genetic marker. Color of blocks indicates genotype or mtDNA haplotype (for the mtDNA) —orange and blue blocks represent homozygous genotype for *P. glaucus* and *P. canadensis* respectively and brown blocks indicate heterozygote genotypes. Black horizontal lines separate individuals by location. (D) Principal Components Analysis showing the individual scores for the first two PCs. The proportion of variance explained by each axis is indicated between brackets. Color indicates whether individuals had the *P. glaucus* (orange), *P. canadensis* (blue), or were heterozygous (brown) for a species diagnostic SNP in *Ldh* and shape indicates an individual’s mtDNA haplotype (*P. glaucus*: diamonds, *P. canadensis*: circles); grey circles were individuals not genotyped for *Ldh*.

The hybrid zone between *P. glaucus* and *P. canadensis* is believed to be maintained primarily by a climate-mediated developmental threshold, produced by geographic variation in the length of the growing season (Scriber & Lederhouse 1992). One line of evidence implicating the growing season is that the hybrid zone coincides with a change in a number of ecologically adaptive traits related to seasonality including the transition of facultative diapause induction of *Papilio glaucus* to the obligate diapause induction of *P. canadensis*. It is presumed that *P. glaucus* alleles involved in facultative diapause would be disadvantageous north of the hybrid zone where on average the growing season is too short to complete two generations (Ritland & Scriber 1985). Conversely, alleles underlying obligate diapause phenotype may be disadvantageous south of the hybrid zone, as they would likely lead to phenotypes with lower fitness than those that emerged for a second generation.

In the *P. glaucus* - *P. canadensis* system, diapause is largely Z-linked (Hagen & Scriber 1989); like birds, Lepidoptera has a ZW system (Z=X, W=Y) of sex determination and females are the heterogametic sex. Divergent selection acting on genes involved in diapause has been invoked as a possible explanation for the observed lack of introgression of mtDNA and Z-linked markers (Scriber *et al*. 2008; Scriber 2011; Scriber *et al*. 2014). Previous studies have documented a dramatic change in allele frequencies of Z-linked allozymes (LDH and PGD), an mtDNA gene (cytochrome oxidase I) and physiological traits (geographic distribution of host-specific detoxification abilities and host plant preferences) that coincide with the environmental gradient that is the length of the growing season (reviewed in Scriber 2011). Given that diapause appears to be an important ecological trait that differs between these species and may be involved in maintaining the location of the hybrid zone, Scriber (2011) suggested that the Z-chromosome may harbor a disproportionate amount of divergence.

In addition to climatic variation, other extrinsic factors may be involved in the maintenance of this hybrid zone, including the distribution of host plants (Lindroth *et al*. 1988; Mercader *et al*. 2009). While both species are highly phytophagous, they differ in their preferences for, and ability to metabolize, linear and angular furanocoumarins, resulting in differential detoxification abilities for some host species (e.g., *Populus tremuloides* can be detoxified by *P. canadensis*, but not *P. glaucus*, with the opposite true for *Liriodendron tulipifera*) (Hung *et al*. 1997). Candidate genes involved in, or linked to, traits associated with various ecological adaptations, such as host plant detoxification, larval chemical defenses, and diapause have been identified (Hagen & Scriber 1989; Li *et al*. 2002, 2003; Cong *et al*. 2015). Yet, how these candidate genes are distributed within the genome and coincide with patterns of genomic divergence, or how they vary with geography and ecological gradients is not well understood. Further, given that extrinsic factors vary in their geographic distribution, they may produce different underlying selective landscapes that could lead to different patterns of introgression across the hybrid zone.

In this study, we characterized genetic differentiation across the genomic and geographic landscape of the ecologically well-studied *P. glaucus* and *P. canadensis* hybrid zone. Specifically, we screened a reduced representation of genetic diversity across the genome using a ddRADseq protocol and characterized genome-wide patterns of differentiation between *P. glaucus* and *P. canadensis* along a 650 km latitudinal transect in the Wisconsin portion of the hybrid zone. First, we set out to characterize genetic structure between the two species and identify regions of the genome potentially impacted by natural selection. We then placed these patterns of differentiation within the genomic landscape to explore the genomic architecture of differentiation and assess whether differentiation is genome-wide or variable among chromosomes (particularly the Z-chromosome). Further, we evaluated the prediction that the distribution of genes previously identified as being putatively involved in ecological adaptations would be associated with regions of the genome under selection and exhibit steep clinal variation across the hybrid zone (i.e., provide further evidence for their being involved in ecological adaptations). Finally, we used a geographic cline approach to explore how significantly differentiated genomic regions (*F_ST_* outliers in allopatry) vary across geography (and exhibit steep clinal variation in sympatry—hybrid zone) and evaluate the relative roles endogenous and exogenous factors may be contributing to patterns of differentiation observed within the genome.

## Methods and materials

### Sample collection and preservation

Three hundred and fourteen specimens were collected between 2007 and 2013 from across a latitudinal transect that spans the *P. canadensis* and *P. glaucus* hybrid zone, from the Upper Peninsula of Michigan through Wisconsin to central Illinois (Table 1; Fig 1A). Of these specimens, all 314 were used to genotype individuals for species diagnostic mitochondrial haplotypes in the cytochrome oxidase subunit1 (*COI*) gene (Sperling 1993), 192 for a species diagnostic Single Nucleotide Polymorphism (SNP) in the lactate dehydrogenase (*Ldh*) gene, and 101 (80 after filtering) used for the generation of reduced representation of the genome-wide diversity using the ddRADseq protocol. Only males were used in all analyses because females cannot be heterozygous for loci on the Z chromosome (females are hemizygous).

**Table 1.** Sampling locations along the transect. The transect is measured as the distance from Mason Co. using the ‘Haversine equation’ and 90° N as longitude and 6,371 km for the radius of the earth.

| Location | County | Latitude | Distance (km) | *n<sub>COI</sub>* | *n<sub>Ldh</sub>* | *n<sub>ddRAD</sub>* |
| --- | --- | --- | --- | --- | --- | --- |
| 1 | Mason | 40.38352 | 0 | 24 | 10 | 11 |
|  | Putnam | 41.22890 | 94 | 1 | - | - |
|  | Rock Island | 41.528373 | 127 | - | - | - |
| 2 | Green | 42.64672 | 252 | 10 | 10 | 2 |
|  | Iowa | 43.05780 | 297 | 39 | 15 | 12 |
|  | Dane | 43.076369 | 299 | - | - | - |
|  | Richland | 43.37849 | 333 | 4 | - | - |
|  | Sauk | 43.42960 | 339 | 11 | 8 | 2 |
| 3 | Columbia | 43.60436 | 358 | 7 | 7 | 2 |
|  | Green Lake | 43.838402 | 384 | - | - | - |
|  | Marquette | 43.89078 | 390 | 5 | 5 | 3 |
|  | Monroe | 43.94569 | 396 | 1 | - | - |
|  | Adams | 44.03663 | 406 | 14 | 13 | 9 |
|  | Juneau | 44.16089 | 420 | 19 | 16 | 5 |
| 4 | Jackson | 44.23612 | 428 | 16 | 15 | 9 |
| 5 | Wood | 44.44880 | 452 | 16 | 15 | 6 |
|  | Portage | 44.47409 | 455 | 1 | 1 | 1 |
|  | Clark | 44.69498 | 479 | 21 | 15 | 8 |
|  | Marathon | 44.89816 | 502 | 7 | - | - |
| 6 | Taylor | 45.16705 | 532 | 30 | 10 | 6 |
|  | Lincoln | 45.36137 | 554 | 19 | 11 | 3 |
| 7 | Oneida | 45.57978 | 578 | 9 | 8 | 1 |
|  | Price | 45.64626 | 585 | 14 | 14 | 10 |
|  | Vilas | 45.95206 | 619 | 32 | 10 | 1 |
| 8 | Iron | 46.26355 | 654 | 5 | - | - |
|  | Ontonagon | 46.51145 | 681 | 9 | 9 | 10 |
| **Total** |  |  |  | 314 | 192 | 101 |

*n<sub>COI</sub>*: # of individuals genotyped for species-diagnostic SNP in *COI* gene *n<sub>Ldh</sub>*: # of individuals genotyped for species-diagnostic SNP in *Ldh* gene *n<sub>RADseq</sub>*: # of individuals sequenced with ddRADseq

### DNA extraction and genotyping/phenotyping of diagnostic markers

Briefly, gDNA was isolated from leg tissue using a phenol:chloroform protocol (Sambrook *et al*. 1989). The two species diagnostic mtDNA haplotypes (*COI* gene) were scored using a Restriction Fragment Length Polymorphism (RFLP) protocol adapted from Putnam *et al*. (2007). A TaqMan SNP genotyping procedure was used to genotype individuals for the Z-linked SNP located in the coding region of *Ldh* (Table S1). This mutation leads to a non-synonymous substitution that is fixed between the two species and is believed to lead to different LDH isozymes (Putnam *et al*. 2007) that may result in thermophysiological differences between the species (see Fig S1). More details on DNA extraction genotyping of diagnostic markers are provided in supplementary material; all data can be found in Data S1.

### ddRADseq library preparation, sequencing, and bioinformatic post-processing

Two reduced-complexity libraries were generated for 101 individuals using a variation of the Restriction-Site Associated DNA fragment (RAD) procedure (adapted from Nosil *et al*. 2012) involving a double digest (ddRADseq; Parchman *et al*. 2012). Briefly, approximately 600 ng of genomic DNA was digested using the restriction enzymes EcoRI and MseI. For each individual, the EcoRI adapter contained a unique 8 to 10 base pairs (bp) index for barcoding. PCR product was then pooled (into a five and 96 individual library), purified with QIAquick Purification Kit (Qiagen), and size selected (400-600 bp range) on a BluePippin System (Sage Science). One library (containing five individuals) was run on an Illumina MiSeq (Notre Dame Genomics Core) and one (containing 96 individuals) was run on an Illumina HiSeq 2000 (BGI, UC Davis) using paired-end sequencing (150 bp reads on MiSeq and 100 bp reads on the HiSeq)

Raw reads were demultiplexed using a custom Perl script and trimmed and cleaned with the program Trimmomatic (v.0.32, Bolger et al. 2014) using default settings. The first 20 bp from all reads was clipped to remove barcodes and cutsite and only reads with a minimum read length of 30 bp were kept. The 3' end of forward unpaired reads were of lower quality and were trimmed to 80 and 60 bp for MiSeq and HiSeq reads respectively. Reads were then aligned to the *Papilio glaucus* genome v1 (Cong *et al*. 2015) using BWA-*aln* (v.0.7.8, Li and Durbin 2009). Read groups were added using custom Perl scripts, soft-clipped using Picard (v.1.33; http://picard.sourceforge.net) and realigned using GATK v.3.3 IndelRealigner (DePristo *et al*. 2011). Variant calling was performed using GATK HaplotypeCaller (McKenna *et al*. 2010; DePristo *et al*. 2011) with the default settings. Filters were applied to all SNPs in the following order. Variants were filtered for bialellic SNPs and indels removed using VCFtools v0.1.12a (Danecek *et al*. 2011). Individuals with excessive missing data (> 90%) were removed (n=3). Rare alleles with a frequency >95% and <5% (pooling all individuals) were removed using VCFtools. GATK’s default “hard filtering” was applied to keep only high quality variants (QD > 2.0, MQ > 30, ReadPosRankSum > −8.0, HaplotypeScore > 13, MappingQualityRankSum >-12.5). Only genotypes with a ~99% likelihood of being correct were kept by removing SNPs with a genotype quality (GQ) < 20. Following these filters, a minimum of 6X coverage for at least 75% of individuals in the “pure” species populations (*P. glaucus*: locations 1 and 2; *P. canadensis*: locations 7 and 8) was then applied (referred to as RAD1 dataset; 28,417 SNPs). For clinal analyses, we used a subset of the RAD1 dataset that had the additional requirement that a SNP had a minimum of 6X coverage in at least 75% of individuals from *all* eight locations. Following this filter, individuals with less than an average of 14X coverage (for all SNPs) were removed (referred to as RAD2 dataset; 18,316 SNPs) because the estimates of heterozygosity for individuals below this cut-off could be biased by low coverage (Fig S2).

### Putative chromosome assignment of genetic markers

To determine the chromosome and orientation of each *P. glaucus* scaffold (while maintaining the orientation of SNPs within the *P. glaucus* scaffold) we used blastx (ncbi-blast-2.2.30+) (Altschul *et al*. 1990; Shiryev *et al*. 2007) to blast all peptide sequences within each scaffold of the *P. glaucus* assembly against the *Bombyx mori* genome using SilkDB v.2.0 (Wang *et al*. 2005). The *B. mori* scaffold with the most significant blast hits (based on Bit scores) was retained and used to determine the chromosome of each *P. glaucus* scaffold and their orientation within a chromosome. Given the high level of synteny between *P. glaucus* and *B. mori* (Cong *et al*. 2015), we expect this approach to reflect a moderately accurate representation of the genomic distribution of our ddRADseq loci.

### Genetic diversity and structure

Genetic structure was assessed using the Bayesian clustering approach of STRUCTURE v2.3.4 (Pritchard *et al*. 2000; Falush *et al*. 2003) using an admixture model with correlated allele frequencies, a burn-in of 100,000 Monte Carlo Markov chains (MCMCs) and a run length of 500,000 iterations, testing clusters (*K*) from 1 to 5; 15 replicates were run for each value of K. We used the RAD2 dataset pruned for linkage disequilibrium (LD) by keeping only1 SNP per 1 kb (n=6,769 SNPs). The results were post-processed using CLUMPAK (Kopelman *et al*. 2015) and the best *K* was chosen following the Evanno’s method (Evanno *et al*. 2005) as implemented in Structure Harvester v.0.6.94 (Dent & vonHoldt 2012).

A Principal Component Analysis (PCA) was also performed in R v 3.2.2 (R Core Team 2013) using the LD-pruned RAD1 dataset and the package *adegenet* v.1.4-2 (Jombart & Ahmed 2011) to provide a complementary visualization of the genetic structure without relying on any model-based assumptions.

We estimated the levels of genetic diversity within each locality from the RAD2 dataset using the per-site nucleotide diversity (π) (Nei & Li 1979), as well as the inbreeding index (*F_IS_*) using VCFtools v0.1.12b. Linkage disequilibrium (LD) was calculated as mean *r*^2^ for all pairwise SNPs within each location using a modified version of plink-1.07 that allows for our large number of “linkage groups” (scaffolds) (Malenfant *et al*. 2015). Confidence intervals were generated by randomly subsampling 100 times seven individuals (smallest n of all locations) from each location. Using these estimates of *r*^2^ we then calculated the distance (in kb) at which *r*^2^ is predicted to fall below 0.2 for each chromosomes (1-28) by fitting the relationship between *r*^2^ and distance between SNPs within each scaffold to a non-linear model (Hill & Weir 1988) describing the rate of LD decay similar to Marroni *et al*. (2011). A subset of the RAD2 dataset, using SNPs with *F_ST_* > 0.9 and filtered to 1 SNP per 1 kb (i.e. if more than one SNP were within < 1 kb within a scaffold, only the first SNP was retained; 228 SNPs) were used to calculate the hybrid index and heterospecific heterozygosity of each individual using the hi.index and int.het functions in the introgress package (Gompert & Buerkle 2010) in the R computing environment (R Core Team 2013).

## Detecting (outlier) loci potentially under selection

We investigated the pattern of interspecific differentiation using two genome scan approaches conducted on the allopatric populations at the extremities of the transect, location 1 and 2 for *P. glaucus* (n = 18) and locations 7 and 8 for *P. canadensis* (n = 18). After removal of monomorphic loci, we first quantified the level of differentiation in allelic frequencies for each SNP using the dataset filtered using just the allopatric populations (RAD1; 28,417 SNPs) with the *F_ST_* estimator of Weir & Cockerham (1984) using VCFtools. We identified SNPs potentially evolving under natural selection using the *F_ST_*-outliers approach originally described in Beaumont and Nichols (1996) and implemented in ARLEQUIN v.3.5 (Excoffier & Lischer 2010). We estimated the 99.9% neutrality envelope of *F_ST_* as a function of the heterozygosity using 500,000 coalescent simulations under a neutral model of population evolution and a Hierarchical Island Model consisting of 10 groups and 100 demes. SNPs above the upper 99.9% quantile were considered under potential divergent selection while those under the lower 0.001% quantile to be under potential balancing selection.

In addition, we used a Bayesian method implemented in BayeScan v.2.1 to estimate the probability that each locus is subject to divergent selection (Foll & Gaggiotti 2008). To account for multiple comparisons and reduce the amount of false positives, we set the prior odds for the neutral model (pr_odds) to 10,000 and used a false discovery rate of 0.01. Twenty pilot runs of 5,000 iterations were used with a thinning interval of 10 and a burn-in of 50,000 iterations followed by 5,000 iterations for output.

Loci were considered as outliers for potential divergent selection if they were identified as significant with both methods (BayeScan and ARLEQUIN). To test whether the outlier loci were non-randomly distributed across chromosomes, a Fisher’s Exact Test was used to compare the distribution of outlier loci for divergent selection and balancing selection to a null distribution, using all linkage groups as well as a separate analysis with just autosomes; a Bonferroni correction was applied to control for multiple testing (α = 0.01/number of linkage groups; alpha = 0.00034 and 0.0038 when using all linkage groups and just autosomes, respectively). This analysis was also done using the “divergence hotspots” (genes that always show larger inter-species variation on both the protein and DNA sequence level) identified in Cong et al. (2015) to determine whether these highly divergent regions were non-randomly distributed across linkage groups and compare their distribution to outlier regions from our study.

### Geographic cline analysis

We evaluated coincidence (amount of congruence among estimates for cline center for each SNP) and concordance (amount of congruence among estimates for cline width for each SNP) using the RAD2 dataset for ddRADseq loci identified as outliers (see above) that were in HWE in each location and filtered to 1 SNP per 1 kb (228 SNPs), as well as mtDNA (*COI*) and Z-linked (*Ldh*) loci. We fitted a combination of equations that describe the sigmoidal shape at the center of each cline and two exponential decay curves (Szymura & Barton 1986, 1991). A total of 15 different models that varied in shape (5 possibilities: having no tails, symmetrical tails, asymmetrical tails, left tail only, or right tail only) and scaling of tails (3 possibilities: using no scale, empirical estimates or best fit) were compared using a Metropolis-Hasting algorithm implemented in the HZAR package in R (Derryberry *et al*. 2014). Posterior distribution of each model parameter was estimated using three independent chains run using 500,000 MCMC steps after a burn-in of 100,000 steps. This process was repeated three times for each model and each MCMC trace was visually inspected to ensure convergence of the results. Model selection was based on the lowest Akaike Information Criterion (AIC) value and parameter estimates of these selected models were used (Akaike 1974). The cline center and width of any two loci were considered non-coincident or discordant if the 2-log-likelihood unit (comparable to 95% confidence intervals) for the upper and lower bound estimate of cline centers or width did not overlap, respectively.

To determine whether the clinal patterns of loci coincide with a climatic gradient, the estimated center of clinal loci were compared to geographic variation in length of the growing season. Growing degree days (GDD) for each county along a transect spanning the hybrid zone were calculated with PRISMs time series datasets modeled using climatologically-aided interpolation at a 4km resolution (PRISM Climate Group 2014). For each year from 1980 to 2012, GDDs were calculated as the sum of the mean daily temperature ((T_min_ -T_min_)/2- T_base_), between March 1st – Oct 31^st^. A T_base_ of 10° C was chosen based on the lower activity threshold of *P. glaucus* estimated by Ritland & Scriber (1985) at 4km pixels in ArcGIS® (ESRI v10.2) and then averaged to the county level. The 32-year mean and standard deviation among years for each county were used to examine the relationship between the total GDDs and distance across this transect. It has been estimated that the GDD requirement for *P. glaucus* to complete two generations is at least 1300 GDDs (a conservative estimate) (Ritland & Scriber 1985). A linear model was fit to the GDD data by location (distance along the transect). The fit from this model was then used to predict the location where 1200, 1300, 1400 GDDs occurs; 1200 and 1400 GDD were used to get an idea of the geographic variance of this climate-mediated threshold. The distribution of other ecological factors—i.e., hosts associated with differential detoxification, *Liriodendron tulipifera*, *Populus tremuloides*, and *Ptelea trifoliata* (Little 1971, 1976) and the Batesian model *Battus philenor* for which *P. glaucus* is a mimic—were also visually compared to the genetic clines (estimated center of the hybrid zone).

## Results

### Summary of ddRADseq data

One lane of Illumina HiSeq 2000 and one MiSeq run produced 346,321,096 and 12,818,490 paired reads, respectively, derived from 101 individuals. After mapping the reads to the *P. glaucus* reference genome, we called 1,835,135 SNPs, of which 28,417 loci showed a minimum of 6X coverage for 75% of individuals from “pure” parental populations and were used for *F_ST_* outlier analysis. Out of these, 18,316 SNPs showed a minimum of 6X for at least 75% individuals for all populations, with an average coverage of 31.1X ± 8sd and average of 0.07% ± 0.6sd missing data across individuals. This second set was used for all other analysis. A detailed summary of sequencing and SNP filtering statistics is provided in supplementary results (Table S2).

### Putative chromosome assignment of genetic markers

We used the close synteny of *Bombyx mori* and *P. glaucus* to infer the linkage group of our ddRADseq SNPs. Synteny is relatively high within Lepidoptera (>80%), and *P. glaucus* and *B. mori* share >85% of their genes in micro-syntenic blocks (Cong *et al*. 2015). For the 28,417 ddRADseq loci used for outlier analysis, 97% mapped to a *B. mori* scaffold, with 92% mapping to a linkage group (Fig S3; Data S2), spanning all 28 *B. mori* chromosomes. As expected if the chromosome assignments were correct, we found a significantly positive relationship (*r*^2^ = 0.69; *P*-value < 0.0001) between *B. mori* chromosome size and number of mapped loci (Fig. S4). Thus, our ddRADseq SNPs appear to be well distributed across the genomes of *P. glaucus* and *P. canadensis*.

### Population structure and diversity

The Bayesian clustering analysis of *STRUCTURE* showed the highest likelihood values and highest rate of likelihood change as the number of clusters (*K*) increased from 1 to 5 was obtained for *K=*2 (Fig. 1B and S5). Most individuals (18 *P. glaucus*, 55 *P. canadensis*) clustered into two distinct groups corresponding to the two species with more than 95% assignment probability to a species cluster (Fig 1B and Data S3). The Principal Component Analysis (PCA) revealed a similar clustering (Fig 1D).

Seven individuals showed an admixed ancestry with less than 95% of their genome coming from either species based on STRUCTURE analysis (Fig 1B, and Fig S5). Of the seven putative hybrids, two appeared to have mixed ancestry resembling early generation hybrids (<60% assignment to either cluster and nearly all diagnostic loci between the two species are heterozygous for both individuals) (see below and Fig. 1C) and five appear to be later-generation hybrids (Fig. 1B, D, and Fig. S6). Five of these putative hybrids were found at sampling location 3, previously identified as the location of the hybrid zone (Scriber *et al*. 2003) where hybrids made up 31% of the individuals from this location. However, using the species diagnostic marker, *Ldh*, for which we have a larger sample size and assuming individuals heterozygous for this marker are early generation hybrids (a reasonable assumption), we get a lower estimate of 15% (6/41) of individuals being putative hybrids in location 3, and only 3% (6/192) across the entire transect.

### Genetic differentiation and detecting outlier loci

Genetic differentiation, measured by mean (weighted) (Weir & Cockerham 1984) *F_ST_* between the two species (RAD1 dataset) was moderate (0.17). However, the distribution of *F_ST_* was slightly bimodal, made up largely of loci with *F_ST_* value <0.25 (89%) and a small subset (3%) with estimates >0.75 (Fig 2C).

**Figure 2:**
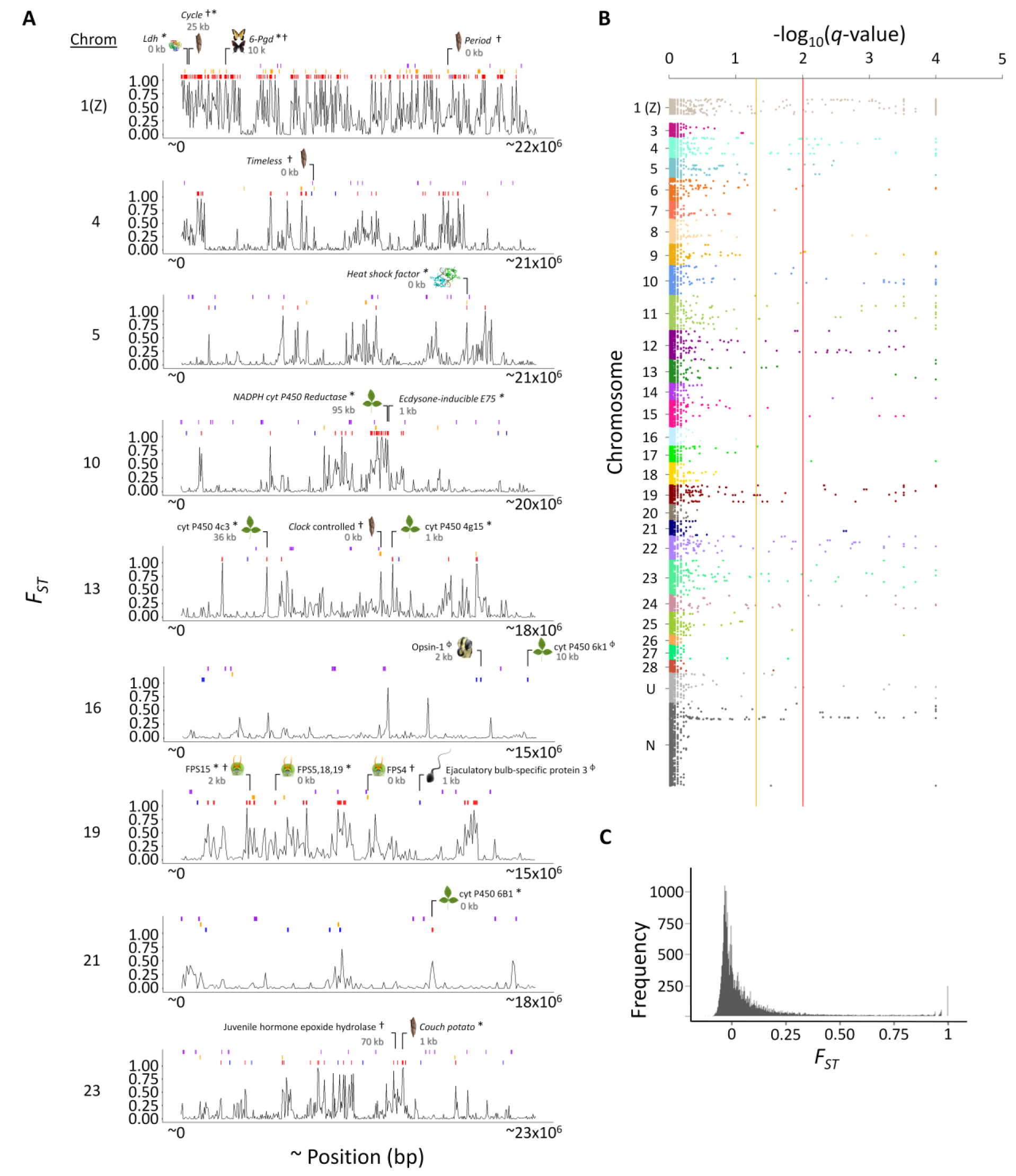
Genomic landscape of divergence between *P. glaucus* and *P. canadensis*. (A) Differentiation (*F_ST_* averaged in non-overlapping 1 kb windows) within linkage groups containing candidate genes for ecologically divergent traits (position of candidate genes shown at the top of each linkage group along with distance from closest ddRADseq locus); note that these distances are less than one cM and the average LD decay of ~80 kb for r^2^ > 0.2. *, ϕ, indicate genes associated with loci under divergent and balancing selection, respectively. ^†^ indicate genes that were identified as divergence hotspots by Cong et al. (2015). Red and blue bars indicate loci under divergent and balancing selection, respectively (*F_ST_*-outlier); orange and purple bars represent loci within 1 kb of genes identified as being divergence hotspots or under positive selection by Cong et al. (2015), respectively. (B) Manhattan plot of log_10_(*Q-values*) from BayeScan analysis by linkage group. Colored lines indicate threshold for False Discovery Rate (orange = 0.05; red = 0.01). Chromosome “U” and “N” represent loci that did not have linkage group information or did not map to a *B. mori* scaffold, respectively. (C) Histogram of *F_ST_* values for all 28,417 ddRADseq loci.

For the 28,417 SNPs we identified, of those that were at least 1 kb apart, 368 (1.3%) were significantly more differentiated in both the *F_ST_*-outlier (Fig. S7A) and BayeScan (Fig. S7B) analysis and could potentially be under divergent selection (Fig 2B). As for those SNPs that were less differentiated than expected under a neutral expectation, we identified 148 (0.5%) falling outside the lower 99.9% envelope (Fig. 2A and S7A). Genetic differentiation between the two species was highly heterogeneous across the genome (Fig. 2A, S8). While most SNPs showed low levels of differentiation (*F_ST_*), a small subset displayed very high *F_ST_* values, with 239 fixed differences (of 28,417 loci), 301 SNPs with *F_ST_* between 0.90 and 0.99 and 234 SNPs with *F_ST_* between 0.80 and 0.89 (Fig 2C). These highly differentiated (*F_ST_* > 0.90) SNPs that were also identified as outliers were distributed across all but five chromosomes (2, 3, 18, 20, and 26) of the *Bombyx* reference genome (Fig 2B, Fig. S8). Some of these chromosomes contained highly divergent regions spanning multiple scaffolds within the chromosome, most notably on the Z-chromosome where regions with high *F_ST_* span nearly the entire chromosome. The Z-chromosome had an average *F_ST_*~4.4 times greater than the genomic background of differentiation compared to autosomes—Z-linked SNPs had an average *F_ST_* of 0.36 ± 0.37 (median= 0.19), and autosomes an average *F_ST_* of 0.08 ± 0.18 (median=0.0006).

Outlier SNPs were not randomly distributed across linkage groups (Fig S3). A significantly greater number of SNPs putatively under divergent selection as well as previously identified divergence hotspots (Cong et al., 2015) mapped to the Z-chromosome (*P*-value < 0.0001) (Fig. 2A, B and S3A). Among the autosomes, six chromosomes (4, 9, 10, 11, 12 and 19) were significantly enriched for outlier loci under divergent selection (*P*-value < 0.001) (Fig S3B). Of these, chromosomes 4, 10, 12 and 19 were also significantly enriched for divergence hotspots. Contrastingly, there were twelve chromosomes (5, 7, 8, 16, 17, 18, 20, 21, 25, 26, 27, and 28) that had significantly fewer outliers than expected (*P*-value < 0.001) (Fig S3A).

### Geographic cline analysis

The prediction that there will be a high degree of coincidence (similar cline center) among genetic and phenotypic clines was highly supported by our data. A majority of the outlier SNPs that were in HWE in each location and filtered to 1 SNP per 1 kb were coincident (81%; 187/230) and concordant (80%; 184/230) with the estimated median cline center of 387 km and 57 km median width (Fig 3B). This estimate places the center of the hybrid zone at ~ 43.8° N (Juneau Co., Wisconsin) (Fig 3B). The estimated center of the hybrid zone (location 3) also coincided with a steep cline in hybrid index and a peak in LD (*r*^2^) and π (Fig 3C). Estimates of LD blocks (distance in bp between SNPs with *r*^2^ > 0.2) were also largest in the hybrid zone (locations 3-5 averaging 75 kb), ranging from 36-78 kb across the transect and averaging 61 kb (se ± 6.5 kb) in size.

**Figure 3:**
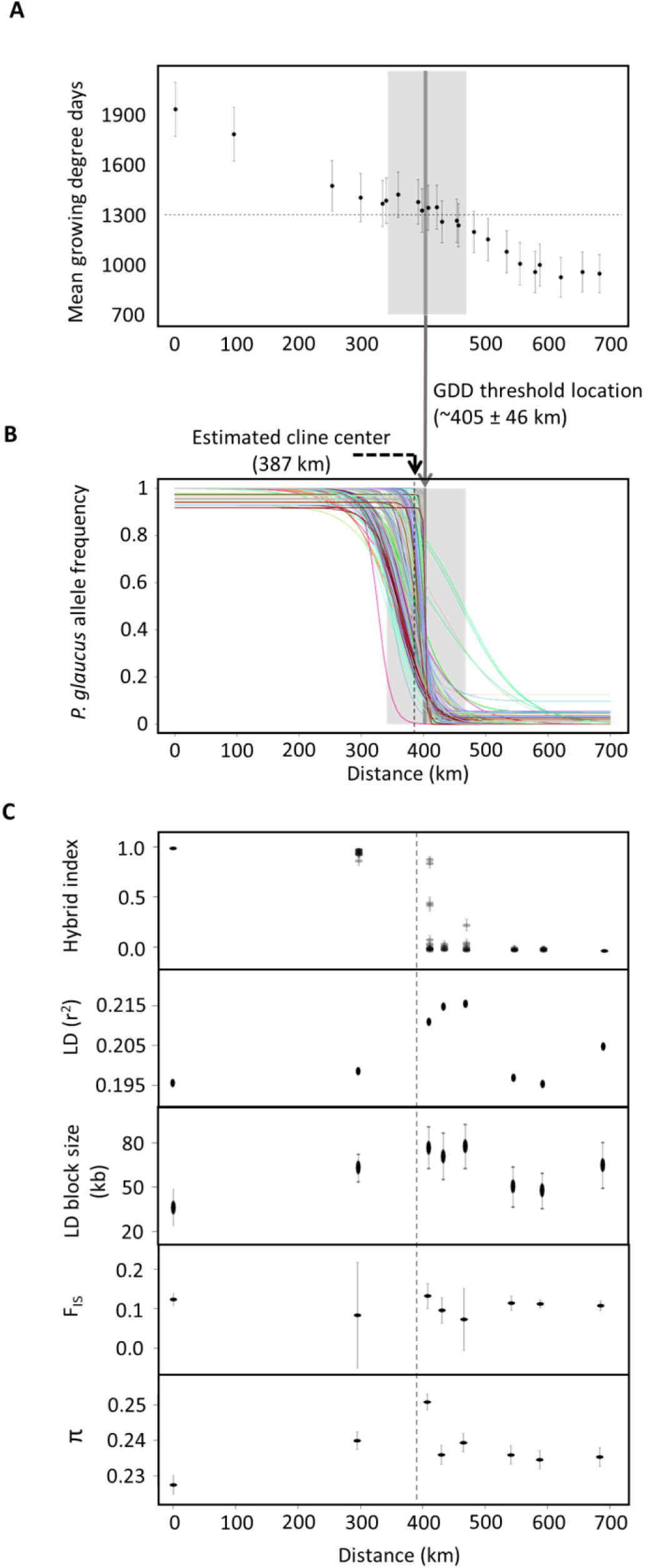
The geographic landscape of divergence. (A) 32 year average length of the growing season (mean growing degree days) across the *P. glaucus* / *P. canadensis* hybrid zone. Error bars are standard deviation. The dark grey shaded region illustrates the (1300) minimum growing degree day requirement (light grey regions ± 100 GDD) for *P. glaucus* to complete two generations (Ritland and Scriber 1985). (B) Maximum likelihood cline for 230 outlier SNPs (in HWE in each location and filtered to 1 SNP per 1 kb). Allele frequency clines that share the same color, map to the same (*B. mori*) chromosome (See Fig 2B). The shaded region illustrates the threshold required for two generations (panel A). (C) Hybrid index, linkage disequilibrium (LD), LD block size (average distance in bp between SNPs with r^2^ > 0.2), fixation index (*F_IS_*) and nucleotide diversity (π) across the *P. glaucus* / *P. canadensis* hybrid zone. Hybrid index ranges from 0 (*P. glaucus-*like) to 1 (*P. canadensis*-like). Error bars represent 95% confidence intervals. Estimates for LD were created by randomly subsampling seven individuals (smallest n of all locations) from each location 100 times. Vertical dashed lines represents the estimated center of the hybrid zone based on genetic clines.

While there was an overall high level of coincidence and concordance across SNPs, 77% of chromosomes (17 of the 22 chromosomes containing SNPs for this analysis) had at least one locus that was discordant or non-coincident (based on non-overlapping confidence intervals with the average estimate) (Data S4). Of the 44 SNPS with non-coincident clines, the majority (40/44; 90%) were shifted in the direction of *P. glaucus*, while only 10% (4/44) were in the direction of *P. canadensis* (Fig 4). SNPs with clines shifting towards *P. glaucus* were found on thirteen different chromosomes. Of the four markers shifting towards *P. canadensis*, two were found on chromosomes 23 while one was on chromosome 1 (Z) and all of these markers were also significantly wider (along with another locus on chromosome 23, they ranked among the top four widest of all clines) (Fig 4).

**Figure 4:**
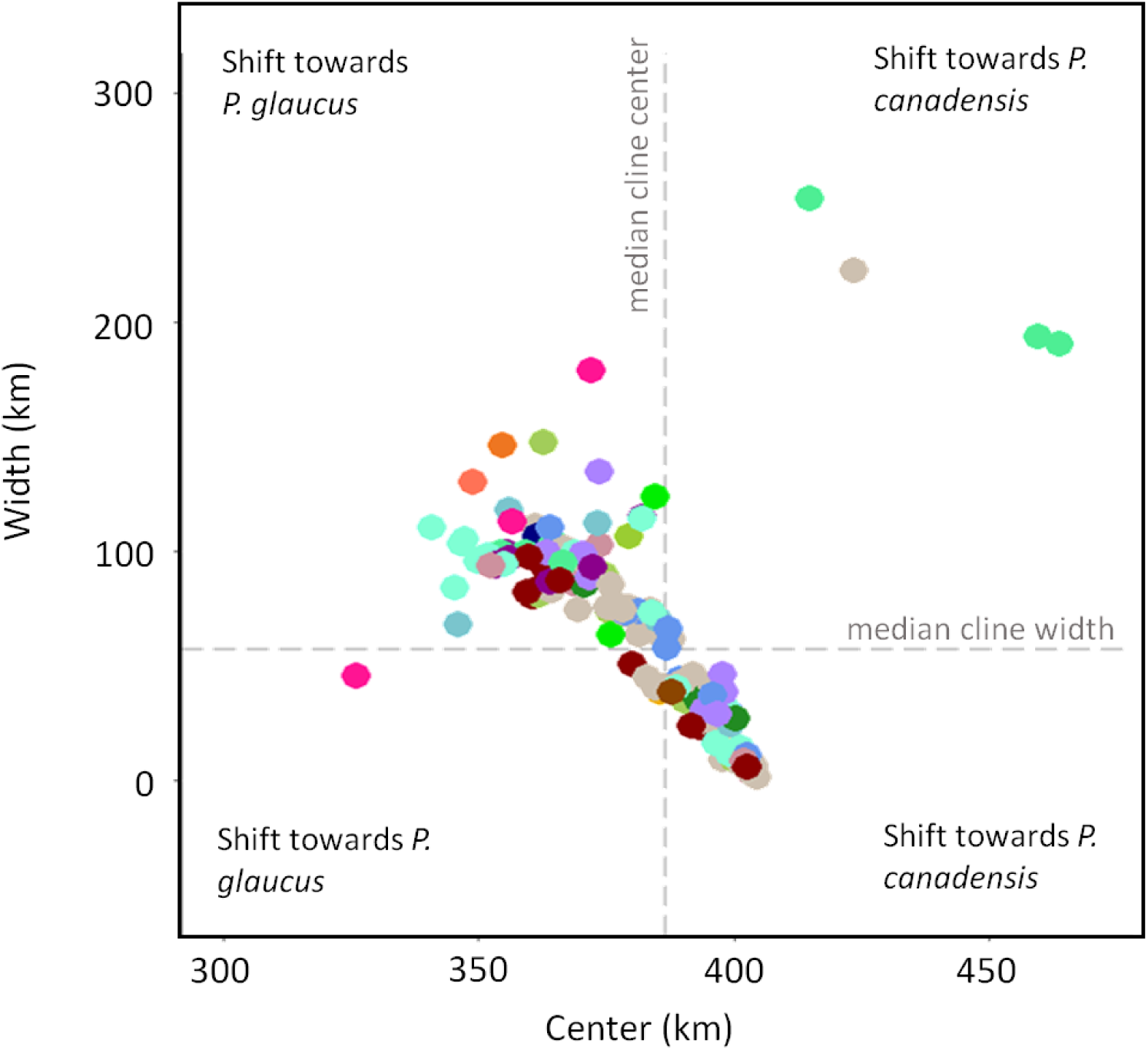
Maximum likelihood estimates of cline center and width for 230 (outlier) SNPs (circles). Dashed lines indicates estimated median cline center (387 km) and width (57 km) based on all genetic markers. Color indicates putative chromosome assignment (see Fig 2B).

Of the only 20% (47/230) of loci that were discordant, 98% (46/47) of these were significantly wider and 2% (1/47) narrower than the median estimate (Data S4). The majority of the significantly wider clines that were also noncoincident (24/28), were shifted to the south of the transect. The only loci found to be significantly narrower than the median cline was a RADseq locus located in a scaffold that did not map to a *B. mori* linkage group. However, it should be noted that given hybrids are rare and make up a small portion of our samples within the hybrid zone, we are only able to detect modest to large differences in cline width among clinal loci.

### Evaluating ecological factors maintaining divergence

The estimated center of clinal loci was compared to geographic variation in length of the growing season. As expected, we found that the boundary of the growing degree day (GDD) requirement to complete two generations coincided with the estimated center of the hybrid zone. The cline center of the genetic loci was 387 km compared with the cline center for the GDD threshold which was at 405 km, putting this climate-mediated developmental threshold ~18 km north of the genetic cline center and similar in cline width (to the genetic loci)—GDD “width” (variance in estimate based on ± 100 GDD) was 64 km, genetic cline width was ~ 57 km (Fig 3B and 4A). Contrastingly, the range boundaries of three host plants (*Liriodendron tulipifera, Populus tremuloides*, *Ptelea trifoliata*), for which differences in detoxification abilities exist between *P. glaucus* and *P. Canadensis*, did not coincide with the hybrid zone. The east-west boundary where *Ptelea trifoliata* reaches its northern limit and *Populus tremuloides* reaches its southern limit is ~100+ kms from the hybrid zone in Wisconsin (Fig 5C, D, E). Similarly, the northern range boundary of the Batesian-mimicry model *Battus philenor* (for the female black morph of *P. glaucus*) does not coincide with the hybrid zone, as its most northern extent ends ~100 kms south of the estimated center of the hybrid zone (Fig 5B).

**Figure 5:**
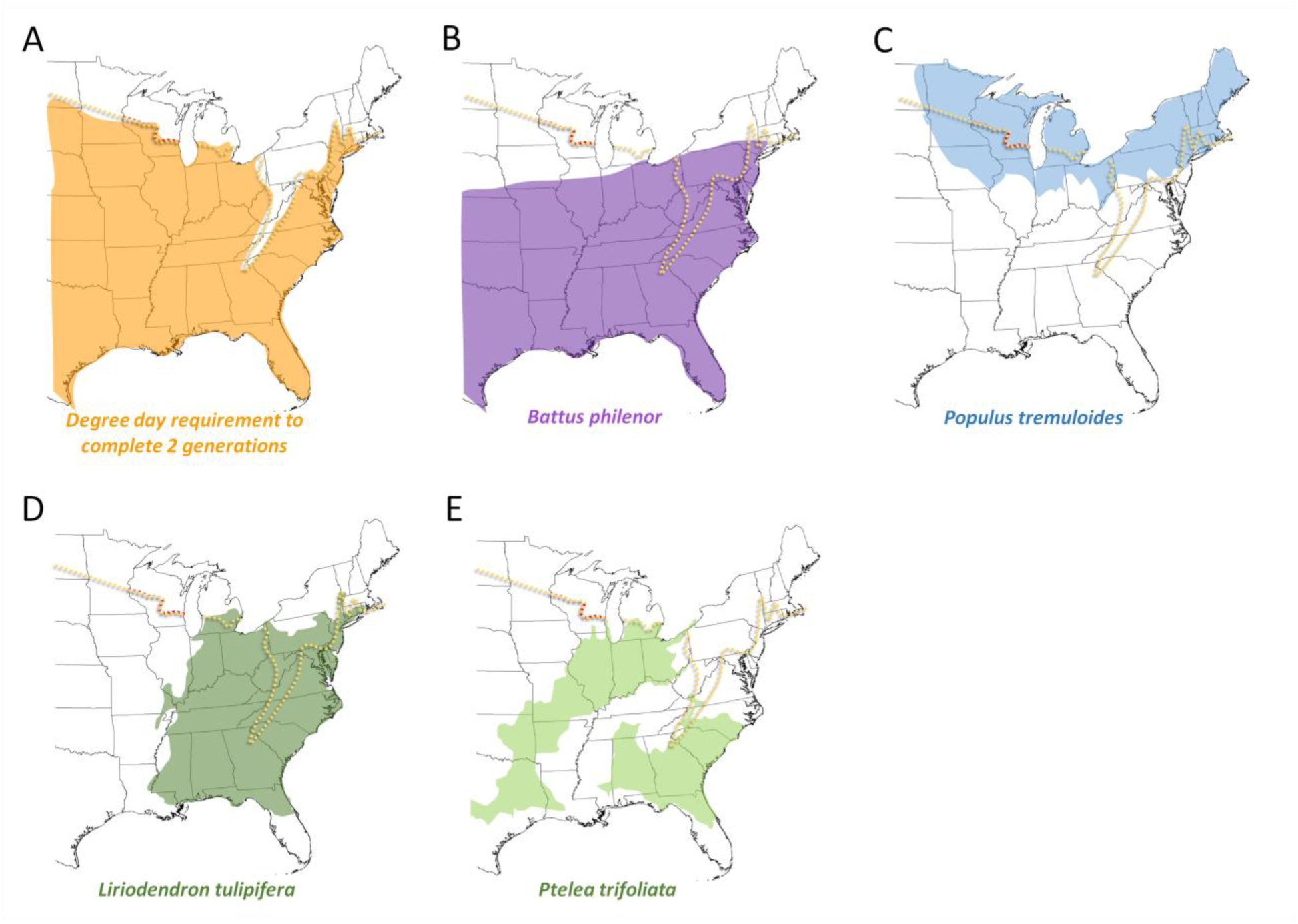
Approximate distributions of ecological factors associated with traits known to be ecological divergent between *P. glaucus* and *P. canadensis* in relation to the estimated location of the hybrid zone. The yellow dotted line indicates the estimated location of hybrid zone and the red portion highlights the region of the hybrid zone sampled in this study. (A) The estimated spatial distribution of growing degree days (GDD) required to complete two generations associated with facultative diapause; adapted from Scriber 2011 a threshold of >1300 GDD. (B) The distribution of the Batesian model (*Battus Philenor*) associated with the female black morph in *P. glaucus*; adapted from previously published distributions of *Battus Philenor* (Fordyce and Nice 2003). (C,D) Estimated distribution of two hosts plants (*Populus tremuloides* and *Liriodendron tulipifera*) associated with differential detoxification abilities between *P. glaucus* and *P. canadensis* (Lindroth *et al*. 1988; Li *et al*. 2004); adapted from range maps published in Little (1971, 1976).

## Discussion

### The genomic landscape of divergence

Placing patterns of divergence within a genomic and geographic context can provide a better understanding of how natural selection and neutral processes drive and maintain species boundaries. Our results, based on a reduced representation of genomic variation, showed that genetic divergence and differentiation between *P. glaucus* and *P. canadensis* is widespread across the genome and is highly heterogeneous in distribution and size within and across chromosomes. A subset of chromosomes (Z-chromosome and the autosomes 4, 10, 12 and 19) showed a disproportionate amount of these differentiated and divergent regions. This pattern of loci putatively under divergent selection distributed across multiple linkage groups, with a sex chromosome harboring what appears to be a disproportionate amount of these divergent regions, is similar to what has been reported in a number of other systems including other butterflies such as *Heliconius* (Martin *et al*. 2013), mosquitoes (Fontaine *et al*. 2015), flies (Tao *et al*. 2003), mice (Teeter *et al*. 2008), and birds (Ellegren *et al*. 2012). This provides further evidence that regions of low recombination, and especially sex chromosomes, may include features that could play a disproportionate role in maintaining or facilitating divergence in the speciation process, as previously suggested (Fontaine *et al*. 2015; Mitchell *et al*. 2015).

Multiple factors can produce variable patterns of differentiation across the genome and interpreting the causes of these patterns depends in part on whether this hybrid zone is a primary contact zone (divergence-with-gene flow) or result of secondary contact (divergence in allopatry), which is not known in this system. However, there are a number of reasons why this hybrid zone is more likely the result of secondary contact. As with many hybrid zones in North America, it is unlikely that this hybrid zone survived climatic fluctuations over the last 0.6 million years of divergence between these two species (Cong *et al*. 2015). More likely, this hybrid zone formed as a result of secondary contact following the last glaciation (Barton and Hewitt, 1985). While not conclusive, the skewed (“F-shape”) *F_ST_* distribution we observe is similar to the pattern observed by Nosil *et al*. (2012) for allopatric *Timema cristinae* stick insect populations, in contrast to the “L-distribution” observed in parapatric *T. cristinae* populations experiencing gene flow. Some models predict that allopatric isolation should result in a more genome-wide distribution of genome divergence (Rieseberg 2001; Noor *et al*. 2001; Nosil *et al*. 2009; Yeaman & Whitlock 2011), in contrast to divergence-with-gene-flow which is predicted to result in a more clustered distribution of divergent loci. Our finding that multiple outlier loci are distributed across all but five chromosomes is thus more consistent with a process of allopatric divergence. However, discordance in the estimated divergence times of Z-linked markers suggests that gene flow has been ongoing during the speciation process (Putnam *et al*. 2007).

Regardless of the geographic origin of divergence, both selection and neutral processes can produce the heterogeneity in differentiation we observe across these genomes. In the case of the Z-chromosome, the nearly chromosome-wide high level of differentiation is likely, in part, the result of the increased genetic drift resulting from the reduced recombination rate and lower effective population size of this chromosome (Oyler-McCance *et al*. 2015). However, there are a few regions that appear to be largely undifferentiated on the Z-chromosome. Whether these regions reflect further variation in recombination within this chromosome, variable gene flow, or are regions that are no longer in synteny (i.e., are actually located on a different chromosomes) is not known. In the case of autosomes, it seems unlikely that variation in recombination alone would explain a pattern that consists of a large number of differentiated regions, distributed widely both within and across nearly all chromosomes. However, there are a number of relatively large differentiated regions that may be the result of suppressed recombination, illustrated by the introgression of what appear to be large genomic blocks (Fig1 C, Fig S8). The importance and potential of inversions in facilitating divergence during speciation has been well documented in a number of systems (Trickett & Butlin 1994; Noor *et al*. 2001; Kirkpatrick & Barton 2006; Hoffmann & Rieseberg 2008; Love *et al*. 2016), however their role in facilitating divergence in this system remains unknown. A recombination and linkage map for *P. glaucus* would help in identifying possible inversions and more generally the role recombination is playing in producing variable patterns of differentiation in this system.

Alternatively, these patterns could reflect variable gene flow. There is now some controversy over whether patterns of heterogeneous differentiation across the genome, assessed by relative measures of differentiation such as *F_ST_*, are caused by variable gene flow or processes that reduce variation in one population (e.g., selective sweeps or variation in recombination rates) (Noor & Bennett 2009; Cruickshank & Hahn 2014). Thus caution should be taken when using these measures alone. However, Cong et al. (2015) previously identified “divergence hotspots” (genes with sequence divergence at both the DNA and protein level that is greater between than within species; regions with elevated absolute divergence), as well as regions under positive selection and found little overlap (11%) between these regions. Similarly, only 10% (36/368) of our ddRADseq SNPs (1 SNP per 1 kb) under divergent selection were closest to a gene identified as being under positive selection by Cong et al. (2015). Although these results are based on a reduced representation of the genetic variation across the genome, this lack of overlap between loci under divergent and positive selection suggest that positive selection is likely only playing a minor role in the variation in differentiation we observe. For variable introgression to be driving this pattern there would have to be ongoing gene flow. Given that all (*F_ST_*) outliers in allopatric populations had relatively narrow clines in the hybrid zone and were largely coincident and concordant suggests the presence of strong reproductive barriers and very little (later generation) hybridization and gene flow. Thus, the observed variation in differentiation across the genome may reflect heterogeneous selection more than heterogeneous gene flow; although both may be playing a factor.

### Evidence of ecological divergence throughout the genome

A major question remaining in speciation biology is how genomic architecture influences divergence (Nosil & Feder 2012). Resolving this question requires an understanding of how genes involved in ecological adaptation or reproductive isolation are arrayed within the genome. We find that candidate genes for divergent ecological traits (seasonal adaptations, host plant detoxification, and predator defense gland) in *P. glaucus – P. canadensis* system are distributed throughout the genome (across at least seven chromosomes), somewhat clumped and almost all associated with regions of elevated differentiation. Loci found to be significantly differentiated in allopatric populations (*F_ST_*-outliers) that also exhibit steep clines across the hybrid zone are strong candidates for reproductive isolation (Harrison & Larson 2016). Interestingly, we find that nearly all ddRADseq outlier loci, spanning multiple chromosomes, exhibit steep clines across the hybrid zone. That the majority of these loci were also highly coincident and concordant, suggests that reproductive isolating factors are spread throughout the genome and/or selection against hybrids is very strong.

While loci involved in reproductive isolation may be distributed across the genome, five chromosomes seem to play a disproportionate role. Chromosomes 1 (Z), 4, 10, 12 and 19 harbor a significantly greater number of loci potentially evolving under divergent selection as well as containing divergence hotspots, and all exhibited steep clinal variation and (with the exception of chromosome 12) contained candidate genes for ecologically divergent traits. The Z-chromosome contains near chromosome-wide elevated differentiation (nearly 4 times the background of autosomes) and is enriched with genes putatively involved in all traits that are known to be ecologically divergent between these species, such as diapause (*cycle* and *period*), thermal tolerance (*Ldh*) and a suppressor gene involved in female mimetic dimorphism (linked to *6-Pgd*), all of which are in, or are near (< 30 kb from) highly differentiated and divergent regions. Reduced recombination of this chromosome is likely facilitating some of this divergence, as a region as large as ~700 kb has been estimated to be in linkage on the Z-chromosome (Winter & Porter 2010). Regions of the genome experiencing low recombination such as near centromeres (Ellegren *et al*. 2012), sites of inversions (Twyford & Friedman 2015) and often sex chromosomes (Sæther *et al*. 2007; Teeter *et al*. 2008) have been shown to accumulate genes involved in adaptive divergence. Both *period* and *cycle* of the Z-chromosome are central to the circadian clock system that regulates the timing of eclosion (Blanchardon *et al*. 2001; Zhu *et al*. 2008) for which Cong et al. (2015) identified clusters of mutations on the surface of these proteins that likely modulate differences in diapause between these species (obligate vs. facultative diapause). Interestingly, while *period* was identified as a divergence hotspot by Cong et al. (2015), we did not identify it as being under divergent selection (based on a number of RADseq loci within 1 kb of this gene). While only 11% of genes under positive selection were also divergence hotspots (as determined in Cong et al., 2015), of the few that were, four were found on the Z-chromosome, and include the gene *period*, suggesting that positive selection, not divergent selection, may be driving divergence in *period*. Thus, positive selection on traits involved in seasonal adaptation(s), including *period*, may be facilitating divergence on this chromosome (Dean *et al*. 2015).

Four autosomes (4, 10, 12 and 19) also stand-out with their enrichment of loci under potential divergent selection (with steep clinal variation) and divergence hotspot genes. Chromosome 10 contains one candidate gene for host plant detoxification (NADPH cytochrome P450 reductase; 95 kb from a RADseq locus under divergent selection) and one associated with development (ecdysone-inducible protein) that clustered closely in the center of this chromosome within what appears to be a large region of differentiation. Surprisingly, while chromosome four contains the gene *timeless*, a gene identified as a divergence hotspot (Cong et al., 2015) involved in seasonal adaptations (diapause induction) (Tauber *et al*. 2007), it was not identified as an *F_ST_*-outlier in our study. In the fruit fly (*Drosophila melanogaster)* and the monarch butterfly (*Danaus plexippus*) this gene is involved in post-translational modifications of the core clock proteins and regulates differences in the incidence of diapause in response to changes in light and temperature (Reppert 2007; Tauber *et al*. 2007). Given that *P. canadensis* has an obligatory diapause and no longer uses photoperiodic cues, the lower than expected (moderate) differentiation we observed for a SNP in this gene may reflect relaxed selection on this trait in *P. canadensis* (Ryan *et al*. 2016).

Interestingly, a number of candidate genes for farnesyl pyrophosphate synthase (FPPS) homologs hypothesized to synthesize terpenes for a predator defense gland (osmeterium) unique to Papilioinidae (Cong et al., 2015) were found on chromosome 19. While the chemistry of this gland is known to be affected by genetic background and varies substantially through larval development, it is not known to what extent the chemical composition of this gland differs between *P. glaucus* and *P. canadensis* (Frankfater *et al*. 2009). We identified at least two FPPS genes in close proximity (< 2 kb) to loci under divergent selection as well as another FPPS gene identified as a divergence hotspot by Cong et al. (2015). Given the large geographic distribution of these species, it is possible that variation in predator communities could be driving divergence in this trait, but this remains to be tested.

Of the few RADseq loci identified as being under balancing selection, three were in close proximity to genes involved in traits that have been implicated as being under divergent or balancing selection in other systems. Two of these SNPs were located on chromosome 16. This includes a SNP within 2 kb of an opsin gene (opsin-1). A recent study of a North American brush-footed butterfly (*Limenitis arthemis*) found clinal variation in a majority of opsin genes, with many also under balancing selection (Frentiu *et al*. 2015). Interestingly the variation in the long-wavelength opsin genes of *<u>Limenitis arthemis</u>* did not affect spectral sensitivity and instead are speculated to play a role in unknown adaptive functions such as mediating responses to temperature or photoperiod. The other SNP on chromosome 16 identified as a candidate for balancing selection was closets (~10 kb) to a cytochrome P450 gene (CYP6k1). Hybrids of *P. glaucus* and *P. canadensis* exhibit detoxification abilities of both parents (Mercader *et al*. 2009). Whether overdominance is driving an increase in diversity of this cytochrome P450 gene (or others) by increasing fitness, for example by expanding an individual’s host breadth, is not known and warrants further investigation. Chromosome 19 contained a SNP identified as a candidate for balancing selection within 1 kb of a gene that’s closest ortholog is the gene EbpIII that is specifically expressed in the ejaculatory bulb and seminal fluid of *D. melanogaster*. Balancing selection has been proposed as an explanation for high allelic diversity in a number of seminal fluid accessory proteins (Acps), proteins that influence reproductive traits such as sperm transfer, sperm storage, female receptivity, ovulation and oogenesis (Chapman 2001). While these are an interesting set of candidate genes for balancing selection in this system, future studies involving more fine scale genomic resolution and functional assays will help to evaluate these candidates more thoroughly.

It should be noted that most of the distances (in bp) between these candidate genes and our RADseq loci are generally less than one cM based on our estimate of a physical-to-map recombination distance of ~322 kb/cM (with the genome being 376 Mb and the total map length 1167 cM). This value is similar to the 180 kb/cM estimated for *H. melpomene* (Jiggins *et al*. 2005). More importantly, these distances were also usually far less than our estimate of the average size of regions in linkage disequilibrium (i.e., those with r^2^ > 0.2), particularly those in the hybrid zone that were estimated to be ~70-80 kb. Moreover, given that regions under selection would likely be at the high end of such estimates, these regions may even be greater than 100 kb in size.

### Climate a salient factor in hybrid zone maintenance

While there are a number of ecologically divergent traits between the two species, climate seems to be a salient factor in maintaining genomic divergence. The first and most prominent reason is that nearly all divergent regions of the genome show a cline pattern that covaries with a climatic gradient—the estimated center of the hybrid zone fell just 20 km south the estimated location of the growing degree day (1300 GDD) threshold to complete two generations; all of the narrowest clines (clines < 5 km in width; those putatively under the strongest selection) are even closer to (~1km south of) this GDD threshold. This estimate is in agreement with previous analyses based on a few species-diagnostic allozymes (Hagen 1990; Scriber *et al*. 2003). Furthermore, while the genetic basis of diapause induction and termination remains elusive (Denlinger 2002), some of the genes most consistently implicated as playing a role in regulating diapause or eclosion phenology—*cycle*, *period*, *timeless, clock—*all mapped to chromosomes identified as harboring a disproportionate amount of divergence. Third, candidate genes for other ecologically divergent traits (i.e., host plant detoxification, and Batesian mimicry) (Zandt Brower 1957; Scriber *et al*. 1996; Mercader *et al*. 2009) do not covary with the ecological gradients that would be associated with these traits, but instead largely covary with a climatic gradient. In the case of three of the host plants associated with the greatest differences in detoxification abilities between *P. glaucus* and *P. canadensis*—*Populus tremuloides*, *Ptelea trifoliata and Liriodendron tulipifera*—the range boundary of these plants fall hundreds of kilometers south of the hybrid zone. Similarly, the northern range boundary of the model *Battus philenor* is located hundreds of kilometers south of the hybrid zone (at least in Wisconsin). Thus, it seems unlikely that the distribution of host plants or a model for Batesian mimicry are playing a major role in determining or maintain the location of this hybrid zone (at least in Wisconsin). Last, there is evidence of steep clinal variation in genetic markers and a number of physiological, phenotypic, and behavioral traits largely coincide with this same climate-mediated developmental threshold in other portions of this hybrid zone (e.g., in Michigan and Vermont) (Scriber 2011).

Examples of climatic gradients driving clines in genetic and phenotypic variation have been well documented across a breadth of taxa, particularly in plants and insects (Knibb *et al*. 1981; Franks & Hoffmann 2012). Specifically, geographic gradients in the length of the growing season that result in changes in voltinism (number of generations in a growing season), so called “voltinism-transition-zones,” often coincide with genetic and phenotypic changes (Bradford & Roff 1995; Ikten *et al*. 2011; Mallet *et al*. 2011). Other examples include adaptation to variation in the length of the growing season that has resulted in clinal variation in the body size of *Pteronemobius spp*. of cricket (Roff 1980), allele frequencies of a circadian gene in the European corn borer (*Ostrinia nubilalis*) (Levy *et al*. 2015), and the critical photoperiod of the pitcher plant mosquito (*Wyeomyia smithii*) (Bradshaw 1976). These examples demonstrate the powerful selective force that variation in the length of the growing season can impose, even in the face of gene flow, given that many of these clines exist within these species’ ranges. The *P. glaucus* and *P. canadensis* hybrid zone also coincides with a voltinism-transition-zone, from bivoltine (facultative diapause of *P. glaucus*) to univoltine (obligate diapause of *P. canadensis*) and it has been hypothesized that strong climate-mediated divergent selection is acting to restrict gene flow, as genes associated with obligate and facultative diapause would be selected against south and north of the hybrid zone respectively (Scriber 2011). Our results lend further support for this hypothesis. Whether, the obligate diapause phenotype (of *P. canadensis*) is an adaptation that predates divergence of these species, or came about post divergence is still not known.

While most clines were largely coincident and concordant, the majority of discordant clines exhibited a pattern of widening (“leaning”) in the direction of *P. glaucus*. This pattern can be an indication of hybrid zone movement (Moran 1981) (i.e., reflect a recent northward shift) or reflect biased gene flow towards *P. glaucus* that could be due to asymmetrical selective pressures or assortative mating (Buggs 2007). Biased introgression into *P. glaucus* would agree with previous estimates of greater gene flow into *P. glaucus* (Zhang *et al*. 2013 and Cong *et al*. 2015). Although introgression appears biased in the direction of *P. glaucus* (southward), the most discordant of all clines were those located on chromosome 23 that were shifted north of the hybrid zone. One explanation of such a pattern could be adaptive introgression (Harrison & Larson 2016). That is, *P. glaucus* alleles are introgressing into *P. canadensis* because they confer some fitness advantage in both genetic backgrounds. Interestingly, two of the SNPs identified as being under divergent selection on this chromosome were within 1 kb of the gene *couch potato*; an amino acid polymorphism in the *couch potato* gene has been implicated in climatic adaptation in *Drosophila melanogaster* by altering reproductive diapause (Schmidt *et al*. 2008). Another SNP was closest to a juvenile hormone expoxide hydrolase, that is implicated in insect development by regulating juvenile hormone (Touhara & Prestwich 1993). Whether these genes are acting to restrict the introgression of favorable alleles on this chromosome is not known.

### Potential for “coupling”

Although climate appears to be a salient factor in the maintenance of this hybrid zone, there are multiple lines of evidence that suggest intrinsic barriers might also be present in this system. One possibility is that the *P. glaucus* – *P. canadensis* hybrid zone is a tension zone that has become associated with a climatic gradient. In support of this idea, we observe a relatively high level of coincidence (81%) and concordance (80%) among both clinal ddRADseq loci that also coincide with a peak in *LD*. These patterns, along with what is estimated as a relatively moderate portion of the genome (1.3%) being under divergent selection have been proposed as an indication of the existence of intrinsic barriers (Bierne *et al*. 2011). Thus it is possible that endogenous barriers have become coupled with an exogenous (climate-mediated) cline to impede hybrid zone movement, due to either lowered population density across this ecotone (Barton & Hewitt 1985) or strong extrinsic selection. Unfortunately, as is the case with many hybrid zones, we do not have robust estimates of population density across the hybrid zone due to the difficulty of obtaining such data. Thus, we cannot completely rule out this possibility. Such patterns do not require invoking endogenous barriers, however, as a geographic-selection-gradient-model can also produce clinal patterns indistinguishable from a tension zone model. Further, the large amount of divergence across the genome may be explained by “genomic hitchhiking” as (extrinsic) selection on many loci, such as that on the candidate genes for ecological adaptations we observe, can result in an increase in genome-wide divergence, even in regions unlinked to known adaptive loci (Feder *et al*. 2012).

Isolation during allopatry can lead to divergence in allelic variation, that when admixed, such as during secondary contact, results in genotypes that are less fit due to the creation of combinations of alleles that were previously hidden from selection (e.g., Dobzhansky-Muller incompatibilities) (Coyne & Orr 2004). If the *P. glaucus* and *P. canadensis* hybrid zone is a secondary contact zone as has been proposed (Scriber *et al*. 1991) and is partly supported by results from this study, it is plausible that neutral process and selection have resulted in the build-up of at least some intrinsic incompatibilities within these genomes. Intriguingly, 40 years of research in this system have produced limited evidence of intrinsic incompatibilities, with reductions in fitness often negligible. Tens-of-thousands of lab pairings from numerous experiments have only observed hybrid offspring exhibiting slightly reduced egg viability (~9%), low-to-moderate pupal mortality (~28%) and a reduction or even increase in fecundity, depending on direction of cross (Scriber *et al*. 1995); there is even some evidence of heterosis, translating into faster developmental rates and larger pupal size (Scriber *et al*. 2003). It is possible that the fitness consequences of hybridization do not manifest until after multiple bouts of hybridization—i.e., in later generation hybrids, such as has been observed in a number of hybridizing species (Bierne *et al*. 2006; Pritchard & Edmands 2013). However, given that we found few early generation hybrids, this suggests that most hybrids are not surviving long enough for multiple bouts of hybridization to occur. Alternatively, it may be that we have been focusing on the wrong phenotype. It has been observed that hybrids of these species exhibit a dramatically altered phenology (emerge weeks to months prior to, or after, parentals) as well as aberrant diapause phenotypes (i.e., failure to diapause even under short day conditions) (Ryan *et al*. 2016). Presumably, these aberrant phenotypes could be less fit regardless of environmental conditions and explain strong selection against admixed genomic backgrounds. Future studies that evaluate the fitness of hybrids and parentals in the field, reared in sympatric (hybrid zone) and allopatric (parental) populations, would help to identify possible intrinsic incompatibilities in this system.

Other non-environmental barriers, such as prezygotic barriers (i.e., mate preferences) could also be preventing hybridization and aiding in the maintenance of this hybrid zone. There is some evidence for mate preference, with what appears to be a preference by males of both species for *P. glaucus* females (Deering & Scriber 2002). However, these preference assays have assumed that males are making the choice, which has never been confirmed and there is actually evidence that females in this system are making the choice when mating (Krebs 1988). Further, hybrids based on our mtDNA and Z-linked species diagnostic markers have a relatively even mix of both *P. glaucus* and *P. canadensis* mothers, suggesting that if there is a preference, it does not appear to be substantial.

The finding that candidate genes putatively associated with host plant detoxification overlap more with a climate-mediated developmental threshold than the geographic distribution of differentially detoxified host plants is intriguing. The simplest explanation for this is that the candidate genes associated with these outlier loci do not actually result in phenotypic divergence in these traits (e.g., differences in detoxification ability). Another possibility is that they are merely linked with other genes under exogenous (climate-mediated) or endogenous selection. This may very well be the case, as the specific CYP450 genes we identified as being under divergent selection are not those implicated in producing the greatest difference in furanocoumarin metabolism. In fact, the RADseq loci closest to CYP6B4 genes (those shown to have the greatest level of furanocoumarin metabolism) exhibited very low levels of differentiation. It is also possible that the CYP6 and CYP4 genes that were implicated as being under divergent selection are associated with detoxification abilities for host plants not explored in this study (*P. glaucus* can feed on plants from > 34 families), or may only play a minor role. An another explanation is that clines for ecological traits not involved in adaptations to climate (e.g., CYP6B and CYP64 genes) have become associated with a climatic gradient as a result of historical cline movement. There is some evidence that tension zones can become “attracted” and move together after contact (Mallet & Barton 1989; Mallet & Turner 1998). Recently Rosser *et al*. (2014) have shown that moving clines can also become trapped when they encounter stationary clines maintained by exogenous selection along an ecotone due to an increase in per locus selection resulting from disequilibrium among clines. Future studies that explore how clinal patterns for a set of genome-wide SNPs vary across the multiple independent portions of this hybrid zone where the distribution of host plants (as well as other ecological gradients) varies should shed light on this question as well as the relative role of endogenous and exogenous barriers in this system (Harrison & Larson 2016).

This study demonstrates the power of assessing how genomic differentiation varies along a gradient of allopatry (pure parental populations) to sympatry (hybrid zone) to reveal insights into the genetic architecture of reproductive isolation. With this approach we show that clinal variation in a climate-mediated developmental threshold can act to maintaining divergence between the genomes of hybridizing species. We also find evidence that previously unrecognized endogenous barriers may be contributing to reproductive isolation in this system, providing some support for the coupling hypothesis that hybrid zones with steep clinal variation associated with an ecotone are the result of a coupling of endogenous and exogenous barriers. Last, similar to what is now a growing number of systems, we find evidence that the Z-chromosome may be playing a disproportionate role in the process of speciation.

## Acknowledgements

Butterfly collections were undertaken by Sean F. Ryan, Mark Scriber, Jason Dzurisin, and Sarah Richman. Many thanks to Benjamin Clifford, Meredith Doellman and Jacqueline Lopez and Melissa Stephens in the Notre Dame Genomics & Bioinformatics Core Facility for their advice in various aspects of experimental design, NGS data collection and analysis. This work was sponsored by NSF grants DEB-0918879 to JMS and DEB-0919147 to JJH and a grant to SFR from the Environmental Change Initiative at the University of Notre Dame.

## Data accessibility

Demultiplexed ddRAD sequence data were deposited in NCBI’s SRA archives under BioProject ID PRJNA388284. Scripts and associated files for processing and analyzing all data in this manuscript were deposited in DRYAD (DOI: http://dx.doi.org/10.5061/dryad.t9221).

## Author Contributions

SFR, JMS, MCF, MEP and JJH designed this research. SFR did field collections and all lab work. SFR performed all analysis, with help from MCF and SO. SO and MCF developed software and scripts for a portion of the genomic analyses. SFR, MCF, JJH, JMS and MEP were involved in writing and editing this manuscript.

